# Robust and Bright Genetically Encoded Fluorescent Markers for Highlighting Structures and Compartments in Mammalian Cells

**DOI:** 10.1101/160374

**Authors:** Anna O. Chertkova, Marieke Mastop, Marten Postma, Nikki van Bommel, Sanne van der Niet, Kevin L. Batenburg, Linda Joosen, Theodorus W.J. Gadella, Yasushi Okada, Joachim Goedhart

## Abstract

To increase our understanding of the inner working of cells, there is a need for specific markers to identify biomolecules, cellular structures and compartments. One type of markers comprises genetically encoded fluorescent probes that are linked with protein domains, peptides and/or signal sequences. These markers are encoded on a plasmid and they allow straightforward, convenient labeling of cultured mammalian cells by introducing the plasmid into the cells. Ideally, the fluorescent marker combines favorable spectroscopic properties (brightness, photostability) with specific labeling of the structure or compartment of interest. Here, we report our ongoing efforts to generate robust and bright genetically encoded fluorescent markers for highlighting structures and compartments in living cells. The plasmids are distributed by addgene: https://www.addgene.org/browse/article/28189953/

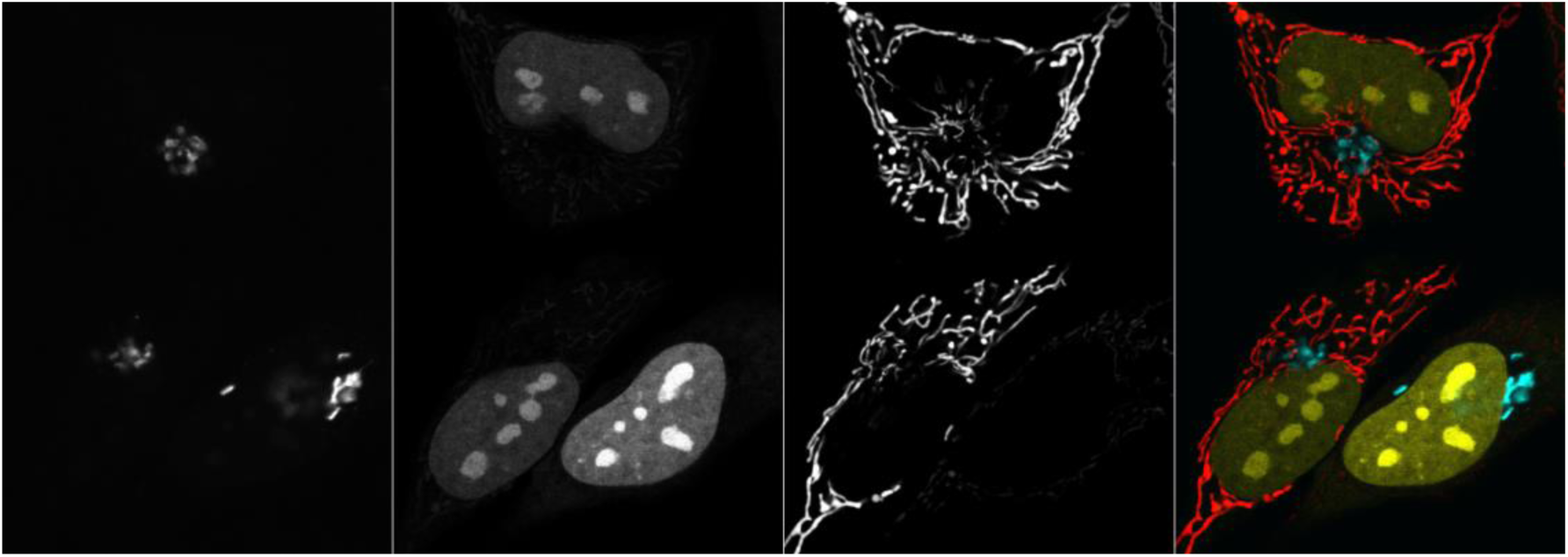

## Introduction

It is difficult to overestimate the impact that Green Fluorescent Protein had and has on basic and applied research. The isolation of the cDNA encoding GFP from the jellyfish *Aequorea victoria* (Prasher et al., 1992) has enabled scientists to track molecules, organelles, protein complexes or cells at different time (milliseconds to days) and length scales (nanometers to centimeters). Fluorescent proteins (FPs) with different emission colors that have since been generated or isolated from different sea-resident animals can be used to highlight independent events simultaneously (Shaner et al., 2005).

By attaching the cDNA encoding a FP to the cDNA of a peptide or protein of interest, specific labeling is achieved. One particular application in cell biology is to use FP fusions as fluorescent markers that highlight structures and compartments in cells. These structures can be visualized with fluorescence microscopy. By employing markers with different emission colors (Rizzuto et al., 1996), the relative location of several structures can be studied (e.g. location of ER relative to mitochondria). Another application of markers is to identify the residency of a biomolecule with an uncharacterized location.

Regardless of the application, it is crucial to use markers that show specific, crisp labeling and minimal spurious, non-specific localization. In addition, it is practical to have a palette of bright fluorescent markers, to facilitate co-imaging. Here, we report our ongoing efforts to generate robust and bright genetically encoded fluorescent markers for highlighting structures and compartments in living cells.

## Results

Previously, we have published a number of markers that highlight a variety of structures and compartments (Bindels et al., 2016; Goedhart et al., 2012). However, we noted that some of these markers also showed spurious localization of the fluorescent protein, i.e. unrelated to the compartment of interest. Below we describe the results of (i) improvements of existing markers and (ii) construction of new combinations of existing markers with established bright fluorescent proteins.

Our choice of FPs is based on (i) high cellular brightness in a spectral class, (ii) good photostability and (iii) monomeric behavior in cells. Based on these criteria, we use the FPs mTurquoise2 (cyan), mNeonGreen (green) and mScarlet-I (red). In the future, this palette of fluorescent proteins may be complemented with other variants, including large Stokes-shift FPs (Shcherbakova et al., 2012) or IR emitting proteins (Shcherbakova et al., 2016) to allow for an even broader range of options.

Before the markers are described, we present an overview of our preferred markers, and their corresponding addgene numbers, in table 1.

**Table 1:** Addgene numbers of the plasmids with genetically encoded markers tagged with mTurquoise2, mNeonGreen or mScarlet-I

| Structure | Targeting sequence | Addgene #<br>(mTurquoise2) | Addgene #<br>(mNeonGreen) | Addgene #<br>(mScarlet-I) |
| --- | --- | --- | --- | --- |
| Nucleus | 3xnls | 98817 | 98875 | 98816 |
| Nucleus | H2A | 36207 | - | 85053 |
| Plasma membrane | Lck (myr/palm) | 98822 | 98878 | 98821 |
| Mitochondria | 4xmts | 98819 | 98876 | 98818 |
| Endoplasmic reticulum | Export&KDEL | 36204 | 137804 | 137805 |
| Nuclear envelope | LaminB | 98830 | - | 98831 |
| Golgi | Giantin | 129630 | 98880 | 85050 |
| Peroxisomes | -SRL | 36203 | 137806 | 85065 |
| Lysosomes | LAMP1 | 98828 | 98882 | 98827 |
| Late Endosomes | Rab7 | 112959 | 129603 | 112955 |
| Actin | ITPKA | 137811 | 98883 | 98829 |
| Actin | Lifeact | 36201 | 98877 | 85056 |
| Actin | UtrCH | 98824 | 98879 | 98823 |
| Microtubules | MapTau | 137807 | 137808 | 112964 |
| Microtubules | α-tubulin | 36202 | - | 85047 |
| Microtubules | EMTB | 137803 | 137802 | 137801 |
| Microtubule tips | EB3 | 98825 | 98881 | 98826 |
| Focal adhesion | Paxillin | 112957 | 129604 | 112956 |

### Nucleus

To target proteins to the nucleus, a strong nuclear localization signal (nls) is usually attached to the protein of interest. As fluorescent proteins are small enough to move out of the nucleus through the nuclear pores, substantial cytoplasmic fluorescence is observed for nls-mTurquoise. To improve nuclear localization, a triple nls, 3xnls (Joosen et al., 2014), sequence was employed. The variant with 3xnls shows increased nuclear labeling relative to cytoplasmic labeling when compared to a fluorescent protein with 1xnls (**Figure 1**). We also noted that the 3xnls variant shows enhanced labeling of nucleoli.

**Figure 1:**
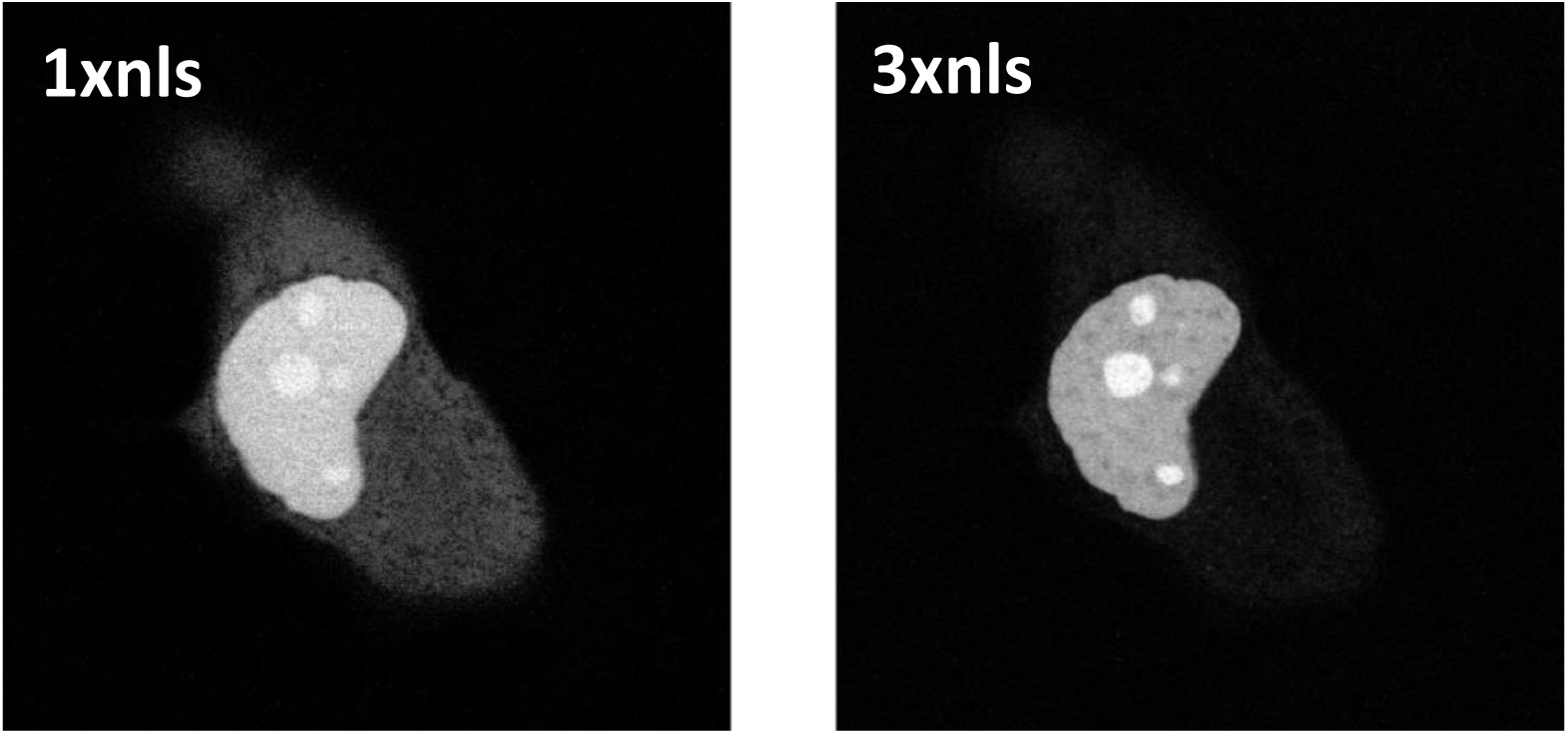
HeLa cell co-transfected with the plasmids encoding a fluorescent protein tagged with 1xnls and 3xnls to highlight the nucleus. The width of the images is 71 µm. (Left: 1xnls-mTurquoise2, right: 3xnls-mScarletI).

The alternative is to use a histone fusion. We have observed that fusions with Histone 2A (H2A, reported here) or histone 2B can be expressed at high levels in cells without noticeable non-specific labeling.

### Plasma membrane

The cysteine-rich sequence of p63RhoGEF was previously used to highlight the plasma membrane (Goedhart et al., 2012). In addition to plasma membrane located fluorescence, we noticed numerous intracellular puncta (**figure 2**, left panel). To create a marker for the plasma membrane with improved selectivity, we explored different peptides. Among the different sequences tested (derived from Lck, Gap43, Src, KRas), we observed the most efficient membrane labeling for the motif from Lck, with hardly any labeling of endomembranes (**figure 2**). Therefore, we recommend to use the Lck-tagged fluorescent proteins as a robust marker for the plasma membrane.

**Figure 2:**
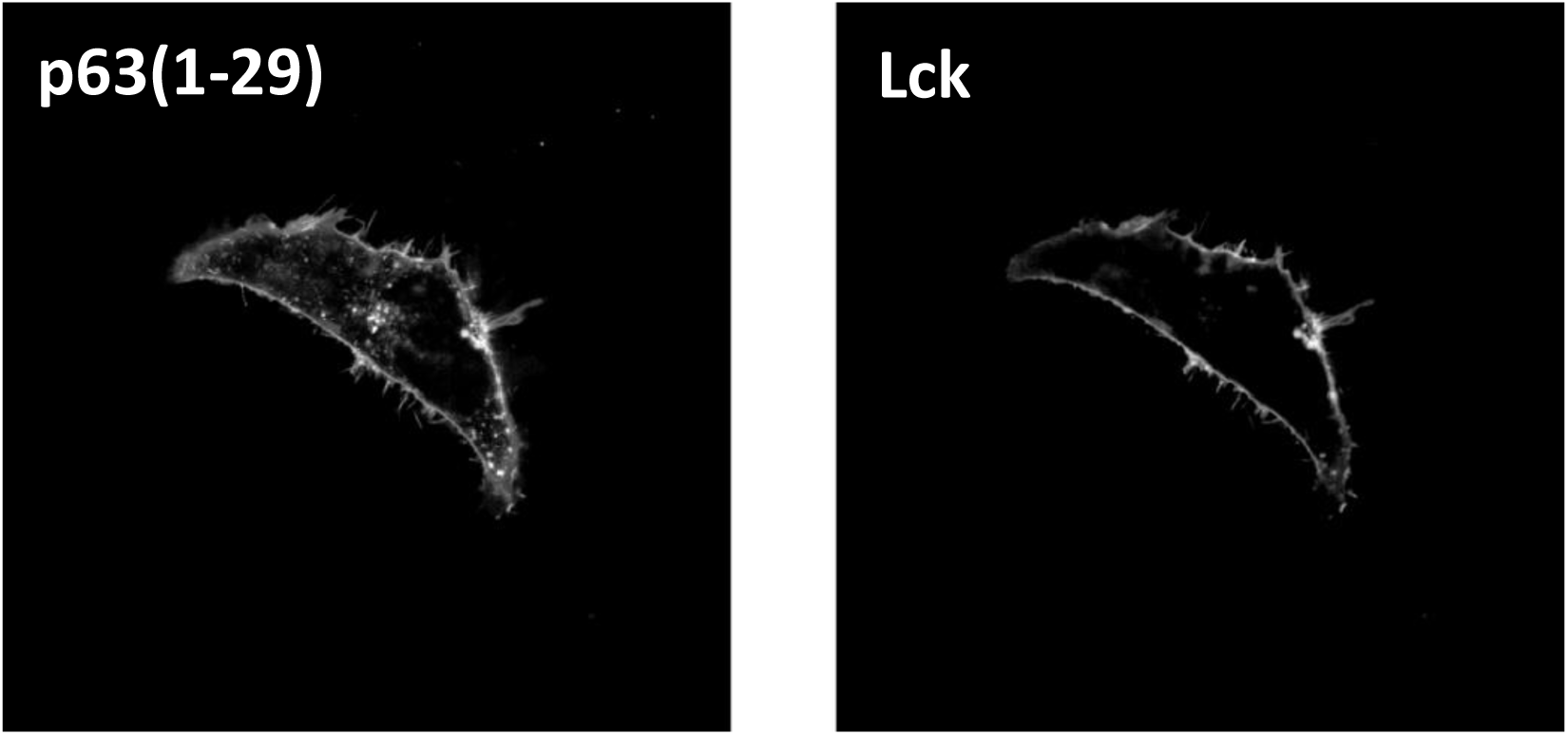
HeLa cell co-transfected with the plasmids encoding a fluorescent protein tagged with the plasma membrane targetin sequence derived from p63(1-29) and Lck. The width of the images is 108 µm. (Left: p63(1-29)-mTurquoise2, right: Lck-mScarletI).

### Mitochondria

We observed that the mitochondrial marker, comprising mTurquoise2 and a signal peptide from COX8A (Goedhart et al., 2012), showed substantial cytoplasmic labeling. To increase mitochondrial targeting, we replaced the single sequence by 4 repeated mitochondrial target sequences (mts) derived from 4mtD3cpv. Co-expression of the 1xmts and 4xmts variant show a crisp labeling of the mitochondria for the 4xmts variant, with hardly any cytoplasmic fluorescence (**Figure 3**).

**Figure 3:**
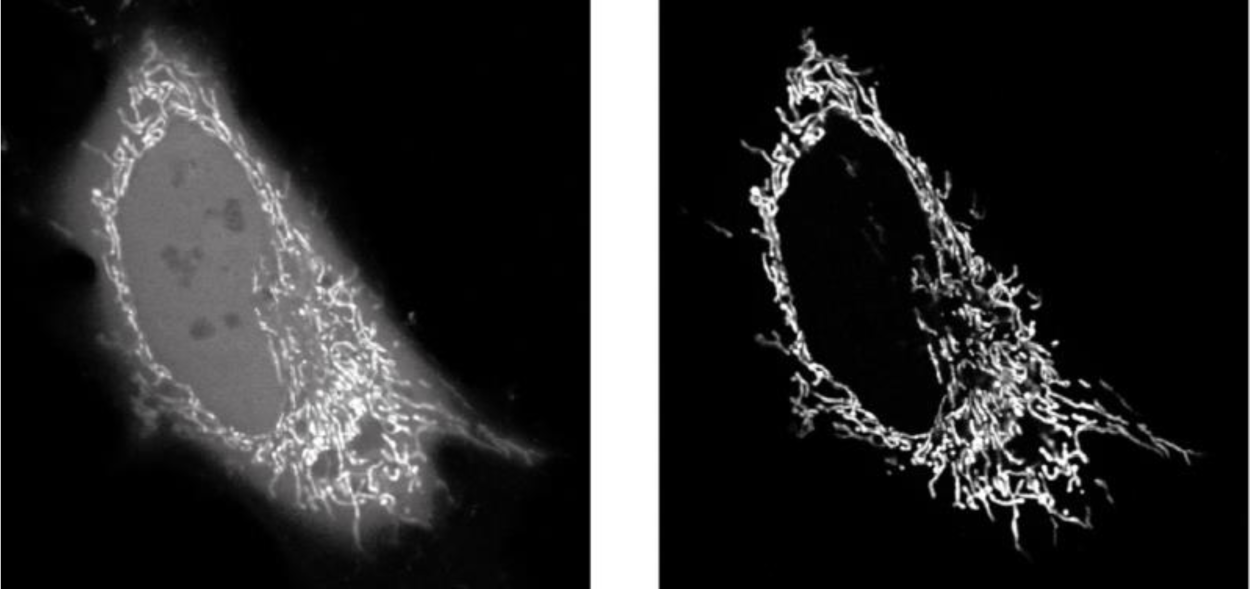
HeLa cell co-transfected with 1xmts-mTurquoise2 (left) and 4xmts-mScarletI (right) to label mitochondria. The width of the images is 71 µm.

### Endoplasmic reticulum and Nuclear Envelope

In our hands, the most robust and crisp labeling of the endoplasmic reticulum in HeLa cells was obtained with a plasmid that was generously provided by Erik Snapp. It uses a signal peptide from chicken lysosome for targeting to the secretory pathway in combination with the KDEL retention signal.

The nuclear envelope is continuous with the endoplasmic reticulum. Still, there are proteins associated with the inner nuclear membrane that can be used to specifically label it. We have examined emerin, lamin A and lamin B and have found that lamin B shows the highest specificity (**Figure 4**). Fusions of full-length lamin B are made to visualize the nuclear envelope.

**Figure 4:**
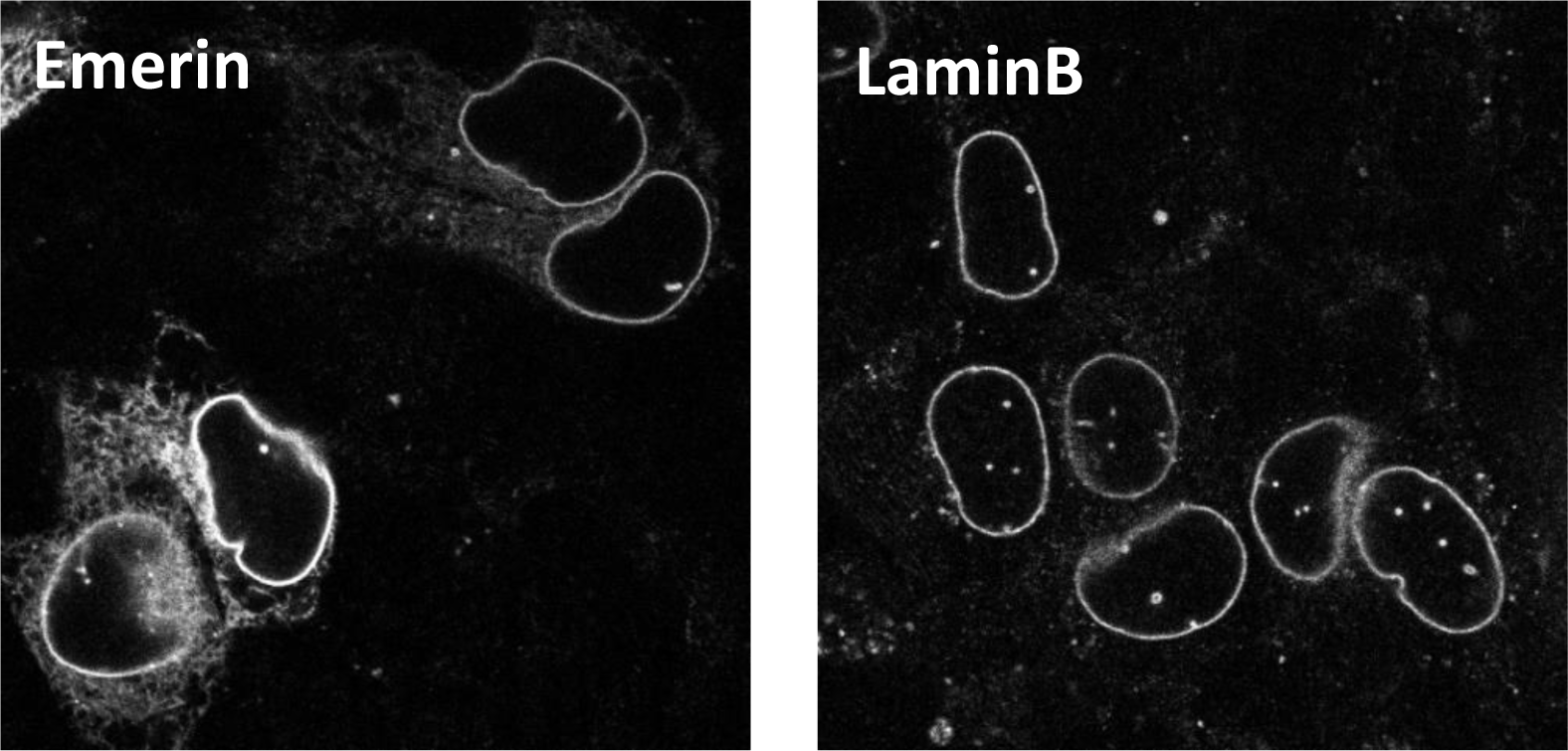
HeLa cells transfected with the plasmids encoding a fluorescent protein tagged with Emerin (left) or Lamin B (right) to mark the nuclear envelope. Width of the images is 73 µm

### Golgi apparatus

Initially, we used a sequence derived from GalT to make a Golgi marker (Goedhart et al., 2012). To evaluate whether this is the optimal marker for Golgi labeling we co-expressed this variant with fluorescent proteins tagged with sequences of Golgi resident proteins, i.e. Giantin (Komatsu et al., 2010) and Sialyltransferase. When introduced into HeLa cells, the fusions with giantin showed labeling that is confined to the Golgi apparatus, whereas the GalT marker showed labeling of punctuate structures, in addition to labeling of the Golgi apparatus (**figure 5**). Therefore, the giantin fusion is the marker of choice for the Golgi apparatus.

**Figure 5:**
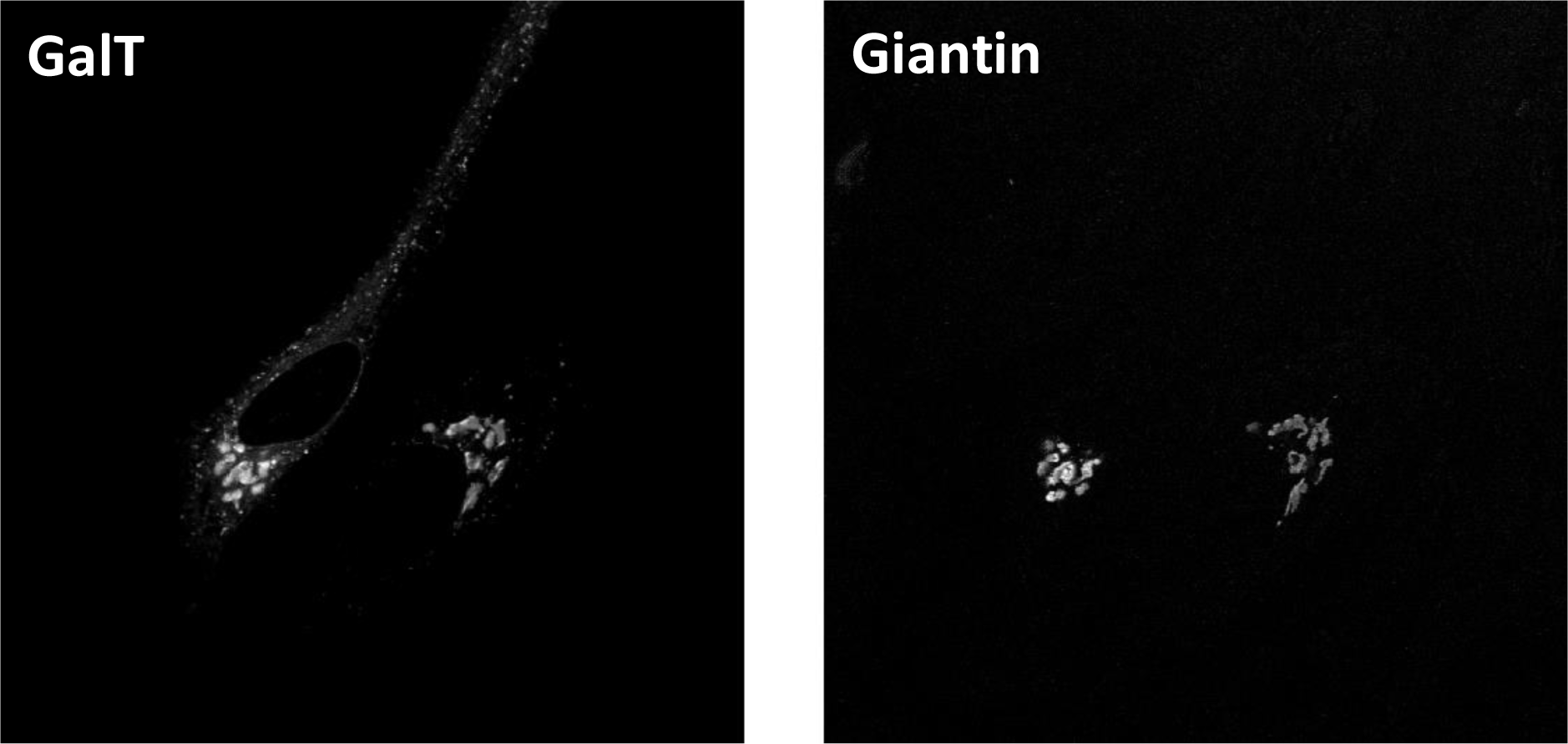
HeLa cell co-transfected with the plasmids encoding a fluorescent protein tagged with GalT and Giantin to mark the Golgi apparatus. The width of the images is 143 µm. (Left: GalT(1-61)-mTurquoise2, right: mScarletI-Giantin(3131-3259)).

### Peroxisomes

Targeting to peroxisomes can be achieved by the c-terminal tripeptide SKL or SRL. Here we have used the SRL signal attached to mTurquoise2, mNeongreen or mScarlet-I to label peroxisomes.

### Endosomes and Lysosomes

For labeling the late endosomes, fusion proteins with full-length Rab7 were made. To construct a lysosomal marker we used the lysosomal associated membrane protein 1 (LAMP1). This fusion protein is expected to decorate lysosomes and indeed, we observed labeling of punctuate structures (**figure 6**)

**Figure 6:**
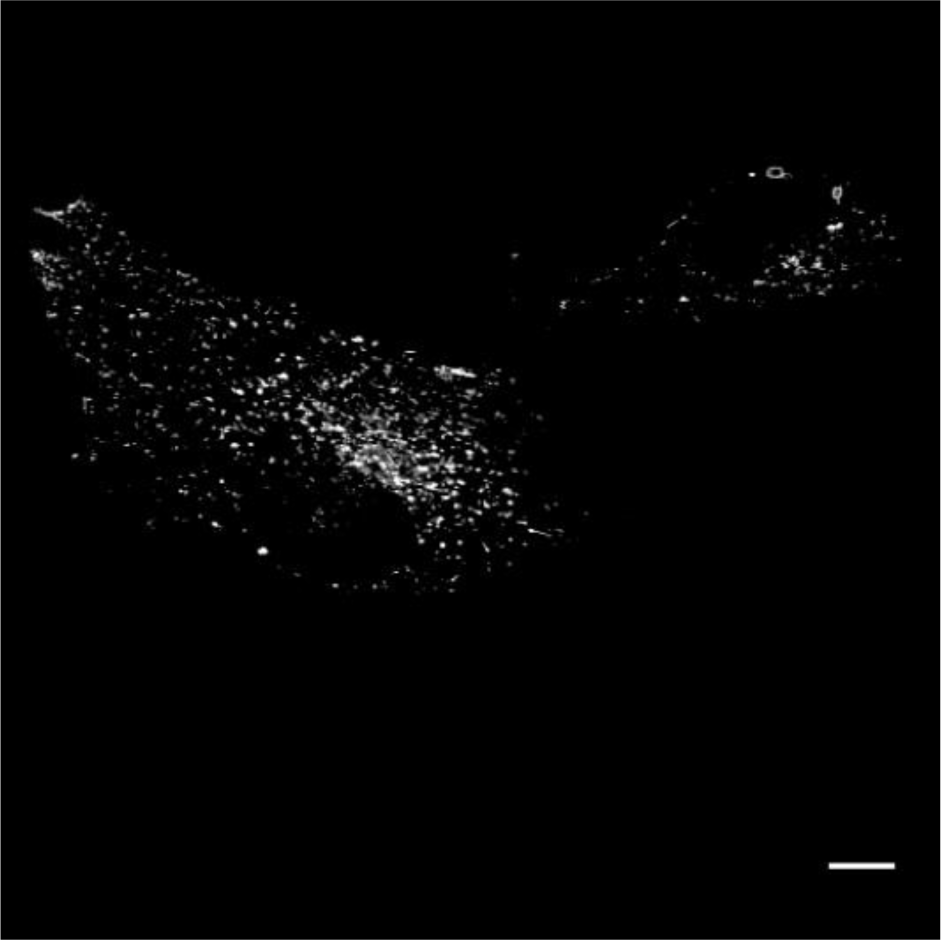
HeLa cell transfected with a plasmid encoding a fluorescent protein tagged with LAMP1 to mark the lysosomes. The width of the images is 143 µm. (mScarletI-LAMP1)

### Actin cytoskeleton

The actin cytoskeleton can be labeled by tagging actin itself with GFP. However, this results in labeling both the soluble, monomeric G-actin pool and the polymerized, filamentous actin (F-actin) pool. To study the cytoskeleton, it is often desired to selectively label actin filaments. To this end, peptide sequences or protein domains have been identified that preferentially bind F-actin, such as lifeact (Riedl et al., 2008), Utrophin-CH (Burkel et al., 2007) and ITPKA(9-52)/F-tractin (Johnson and Schell, 2009). Previously, we have described lifeact fusions with mTurquoise2 (Goedhart et al., 2012) or mScarlet (Bindels et al., 2016). To allow for a wider choice of markers, we made combinations of the bright fluorescent proteins mTurquoise2, mNeonGreen and mScarlet-I with the actin probes Lifeact, Utrophin-CH and ITPKA (residue 9-40). Note that we used a shorter version of Ftractin here (the original Ftractin probes corresponds with residue 9-52 from rat ITPKA, while here we use residue 9-40), without any noticeable difference. All three fusion constructs show pronounced labeling of the actin cytoskeleton (**figure 7**). It is of note that different markers may label different F-actin pools and may have different properties (Belin et al., 2014; Ganguly et al., 2015; Spracklen et al., 2014). Therefore, validation in the system of choice is warranted and the choice of the marker depends on the application.

**Figure 7:**
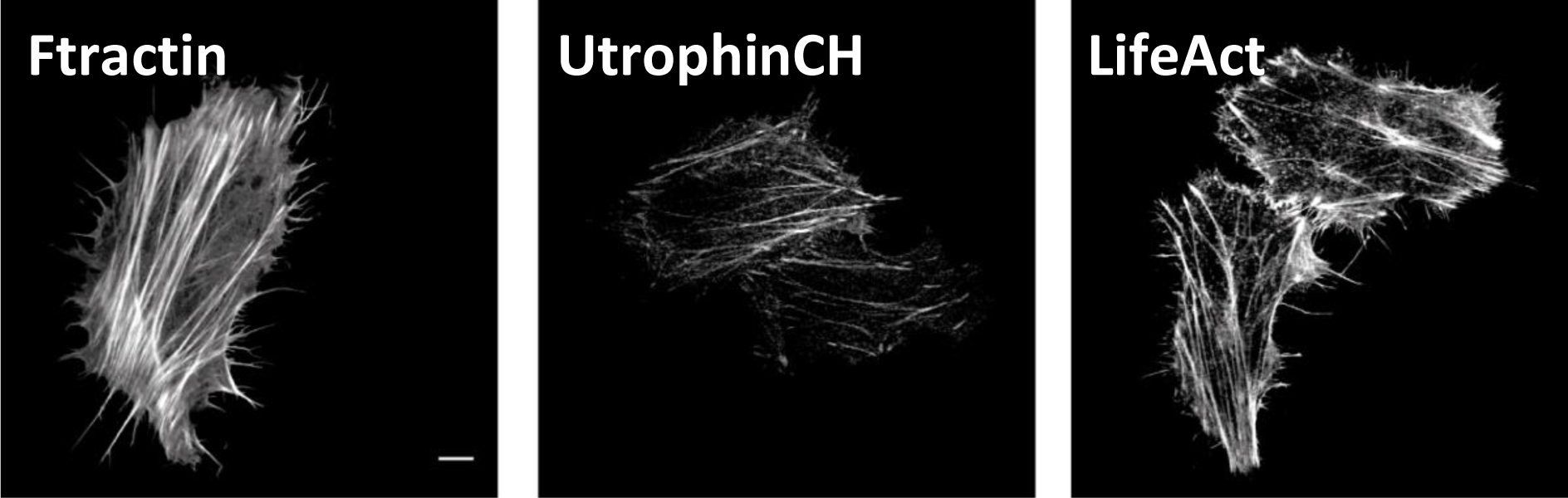
HeLa cell transfected with a plasmid encoding a fluorescent protein tagged with ITPKA or UtrophinCH or Lifeact to visualize the F-actin cytoskeleton. The width of the images from left to right is 116 µm, 143 µm and 143 µm. (From left to right: ITPKA(9-40)-mTurquoise2, mNeonGreen-UtrCH, Lifeact-mScarletI).

### Microtubules

To label the microtubules, FPs can be fused either to the N-terminus of alpha-tubulin (Bindels et al., 2016; Goedhart et al., 2012) or to the C-terminus of beta-tubulin. The protein level of fusion tubulin should be carefully controlled for optimal results. Too much expression leads to higher cytoplasmic signals from unpolymerized tubulin, and also affects the microtubule dynamics. Detailed analyses on the effects by tubulin fusion protein expression are reported with fission yeast (Snaith et al., 2010).

To indirectly label the microtubules, we have generated fusion with the microtubule binding domain of microtubule associated proteins (MAPs) such as EMTB and MapTau (**Figure 8**). Although these are highly efficient markers for microtubules, care should be taken since these proteins may facilitate stabilization and bundling of microtubules.

**Figure 8:**
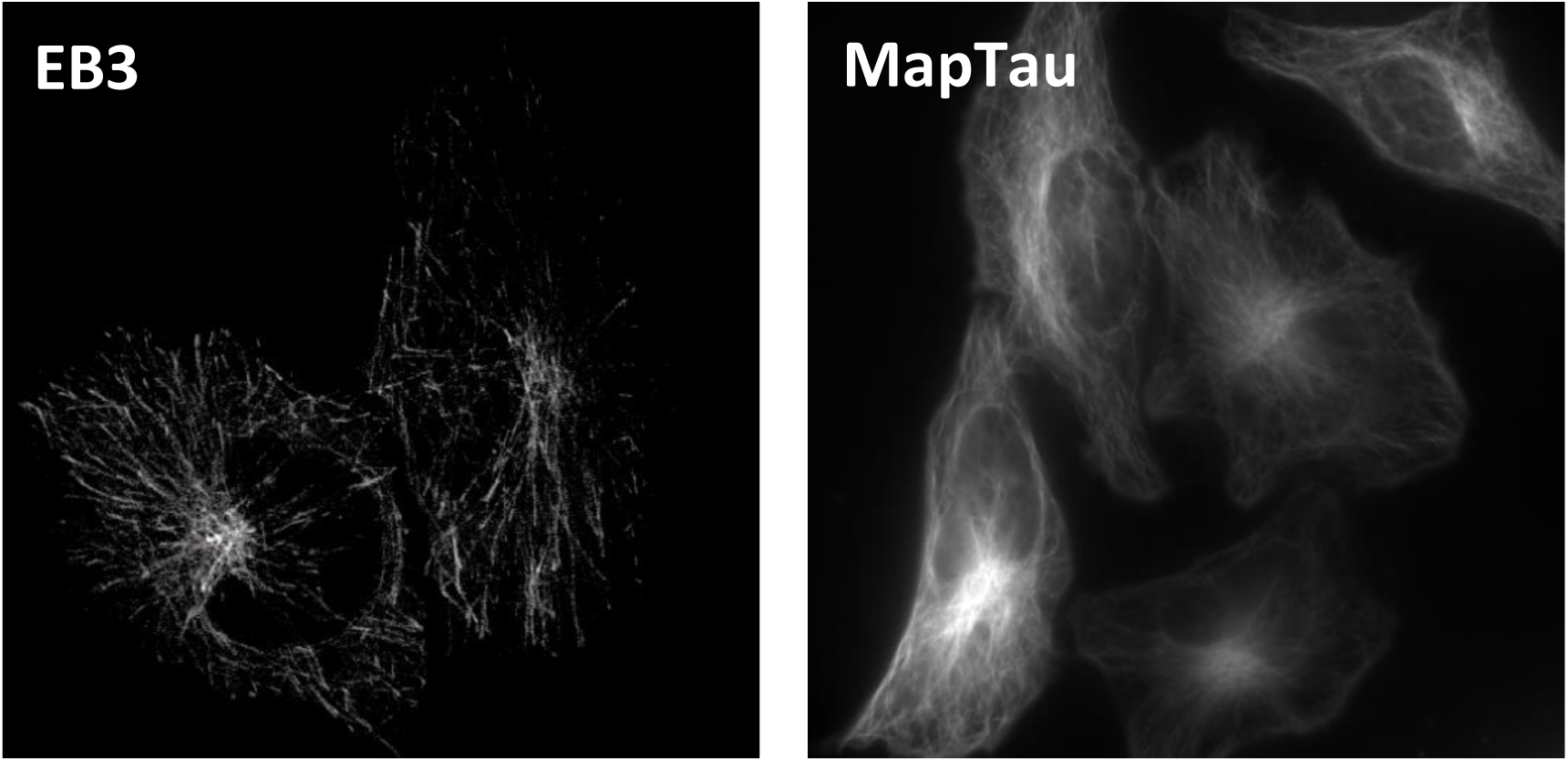
HeLa cell transfected with a plasmid encoding a fluorescent protein tagged with EB3 to detect the growing tips of microtubuli (EB3-mTurquoise2) or with MapTau to label the entire microtubules (mTurquoise2-MapTau).

Another class of microtubule binding proteins, +-TIPs that bind preferentially to the growing plus-ends of the microtubules can be used as the marker for microtubule growth (Stepanova et al., 2003). Among them, EB1 or EB3 fused with an FP is widely used. We observed “comet”-like staining of the growing ends of microtubules with both EB1 (not shown) and EB3 based fusion constructs of mTurquoise2 (**Figure 8**), mNeonGreen, and mScarlet-I. Their brightness and the photostability along with the rapid exchange of the bound populations with the cytoplasmic pool enabled live imaging of microtubule growth at sufficiently low expression levels. A low concentration of the marker is essential for correct measurement of the microtubule dynamics. Higher expression levels can cause hyperstabilization of the microtubules, lower contrast to the background signals from the cytoplasmic pool. At even higher expression levels, their signals extend from the tips to the body of the microtubules.

### Focal adhesions

Focal adhesions are large multiprotein assemblies. Several proteins specifically accumulate in the focal adhesions and these can be used as markers. Thus far, we have only generated fusions with a typical focal adhesion protein, paxillin. It should be noted that this marker shows a considerable fraction of non-bound protein, which may occlude the labeling of the focal adhesion. Lowering the expression level or using total internal reflection microscopy may improve the quality of the labeling.

## Conclusion

Fluorescent markers are essential components of the cell biologist’s toolkit. Several of the previously reported markers show non-specific, suboptimal labeling of structures or compartments. Here, we report markers with improved labeling of the structure or compartment of interest with reduced non-specifc labeling. In addition, we have generated new combinations of markers with the brightest monomeric fluorescent proteins currently available. Our analysis of the labeling patterns and improvements is limited to observations after transient transfection in HeLa cells. It is highly recommended that independent verification of the labeling quality is performed. This is of particular importance, if the markers are intended for different cell lines or organisms. To conclude, we express our hope that the improved markers will be a valuable addition to the tools that are available for cell biology.

## Materials and Methods

### Plasmids

The bright monomeric fluorescent proteins used in this study, mTurquoise2 (Goedhart et al., 2012), mNeonGreen (Shaner et al., 2013) and mScarlet-I (Bindels et al., 2016) have been described previously. Plasmids with Lck (van Unen et al., 2016) Giantin (Bindels et al., 2016; van Unen et al., 2015) and H2A (Goedhart et al., 2012) have been described before.

Plasmids with mCherry-LaminB (Bas van Steensel), EB3-EGFP-N1 (Niels Galjart) EGFP-LAMP1, EGFP-Rab7 (Eric Reits), ER-tdTomato (Erik Snapp), YFP-SRL (Joop Vermeer) were kind gifts.

Plasmids with pcDNA-4mtD3cpv (addgene Plasmid #36324), GFP-UtrCH (addgene plasmid #26737), 3xnls (addgene plasmid #60492), MapTau (addgene plasmid #55077), EMTB-3xGFP (addgene plasmid #26741), YFP-Paxillin (addgene plasmid #50543) were from www.addgene.org

In most cases, existing plasmids were used to generate new combinations by exchanging the fluorescent protein in the clontech-style plasmid that encodes the marker (usually using the restriction enzymes AgeI/BsrGI), with the following exceptions:

The 4mts-mTurquoise2 is made by a partial digest of pcDNA-4mtD3cpv (Addgene Plasmid #36324) with HindIII and the insert is ligated in mTurquoise2 cut with HindIII.

The Utrophin CH domain is excised from the plasmid GFP-UtrCH (addgene plasmid #26737) with BsrGI and SacI and ligated in mTurquoise2-C1 cut with BsrGI and SacI.

LAMP1-mTurquoise2 was generated by a PCR on LAMP1-GFP (primer sequences: forward, 5’-AGCGTCGACATGGCGGCCCCCG-3’ and reverse, 5’ AGCGGATCCTTGATAGTCTGGTAGCCTGCGT-3’), digestion SalI and BamHI and ligation in the pmTurquoise2-N1 plasmid, cut with the same enzymes.

ER-mTurquoise2 uses aa 1-31 of chicken lysosome as export sequence and KDEL as an ER retention signal

The EMTB microtubule binding domain was excised from the plasmid EMTB-3XGFP (addgene plasmid #26741) with AgeI and EcoRI and ligated in mScarlet-I-N1 cut with AgeI and EcoRI.

The coding sequence for YFP in the plasmid YFP-paxillin (addgene plasmid #50543) was excised with AgeI and BglII and replaced by mTurquoise2 that was cut from pmTurquoise2-C1 with AgeI and BglII.

Full plasmid sequence information is available at https://www.addgene.org/browse/article/28189953/ and can be directly accessed through the hyperlinks connected with the addgene numbers that are listed in table 1.

### Cell culture & Sample Preparation

HeLa cells (American Tissue Culture Collection: Manassas, VA, USA) were cultured using Dulbecco’s Modified Eagle Medium (DMEM) supplied with Glutamax, 10% FBS (without antibiotics). Cell culture, transfection and live cell microscopy conditions were previously described (REF). Transfection was performed with lipofectamine 2000, using 1-2 µl of lipofectamine, 50 µl of OptiMEM and ∼500 ng of plasmid DNA per 24 mm coverslip. Alternatively, we used PEI, with 4.5 µl of PEI, 200 µl of OptiMEM and ∼ 500 ng of plasmid DNA per 24 mm coverslip.

### Microscopy

Images were acquired on a Zeiss LSM META confocal microscope with settings as follows. For mTurquoise2 we used 458 nm excitation and a BP480-520 nm emission filter. For mNeonGreen we used 488 nm excitation and a BP500-530 nm emission filter. For mScarletI we used 543 nm excitation and a LP560 nm. Alternatively, we use a Nikon A1 confocal microscope with settings as follows. For mTurquoise2 we used 458 nm excitation and a 482/35 nm emission filter. For mScarlet-I we used 561 nm excitation and a BP 595/50 nm emission filter. All images were acquired with the pinhole set to 1 airy unit.

## Availability of materials

Plasmid DNA encoding the markers described in this study will be distributed by Addgene, together with full plasmid sequences and additional information at https://www.addgene.org/browse/article/28189953/

## Acknowledgments

The cDNAs encoding mCherry-LaminB (Bas van Steensel), EB3-EGFP (Niels Galjart) EGFP-LAMP1, EGFP-Rab7 (Eric Reits), ER-tdTomato (Erik Snapp, Albert Einstein College of Medicine), YFP-SRL (Joop Vermeer) were kind gifts from colleagues.

## Competing interests

The authors declare no competing or financial interests.

## Author contributions

All authors participated in this study, either by generating plasmids, acquiring fluorescence images or writing (or a combination thereof).

## Supplemental information

### Amino acid sequences of the targeting peptides or protein domains

**1xmts** is residue 1-29 of HsCOX8A:

MSVLTPLLLRGLTGSARRLPVPRAKIHSL

**4xmts:**

MSVLTPLLLRGLTGSARRLPVPRAKIHSLGDPMSVLTPLLLRGLTGSARRLPVPRAKIHSLGKLATMSVLTPLLLRGLTGSARRLPVPRAKIHSLGDPMSVLTPLLLRGLTGSARRLPVPRAKIHSLG.

**1xnls** is residue 126-132 of the SV40 large T antigen:

PKKKRKV

**3xnls** is:

MGLRSRADPKKKRKVDPKKKRKVDPKKKRKV

**Lck** is residue 1-10 of MmLck:

MGCVCSSNPE

**p63(1-29)** from Hsp63RhoGEF is:

MRGGHKGGRCACPRVIRKVLAKCGCCFAR

**GalT** has residue 1-61 of HsB4GALT:

MRLREPLLSGSAAMPGASLQRACRLLVAVCALHLGVTLVYYLAGRDLSRLPQLVGVSTPLQ

**Giantin** is residue 3131-3259 of HsGiantin:

EPQQSFSEAQQQLCNTRQEVNELRKLLEEERDQRVAAENALSVAEEQIRRLEHSEWDSSRTPIIGSCGTQEQALLIDLTSNSCRRTRSGVGWKRVLRSLCHSRTRVPLLAAIYFLMIHVLLILCFTGHL

**IPTKA** also known as **Ftractin** is residue 9-40 of Rat IPTKA:

MGMARPRGAGPCSPGLERAPRRSVGELRLLFEA

**UtrCH** is residue 1-261 from HsUtrophin

**Lifeact** is residue 1-17 from Abp140 (Saccharomyces cerevisiae):

MGVADLIKKFESISKEE

**Peroxisomal targeting signal** is -SRL.

**ER** is residue 1-31 of chicken lysozyme preceding the fluorescence protein: MRSLLILVLCFLPLAALGKVFGRCELAAAMK and KDEL at the C-terminus of the fluorescent protein.

## Notes

https://www.addgene.org/browse/article/28189953/

